# Cryo-EM structure of haemoglobin at 3.2 Å determined with the Volta phase plate

**DOI:** 10.1101/087841

**Authors:** Maryam Khoshouei, Mazdak Radjainia, Wolfgang Baumeister, Radostin Danev

## Abstract

With the advent of direct electron detectors, the perspectives of cryo-electron microscopy (cryo-EM) have changed in a profound way^1^. These cameras are superior to previous detectors in coping with the intrinsically low contrast of radiation-sensitive organic materials embedded in amorphous ice, and so they have enabled the structure determination of several macromolecular assemblies to atomic or near-atomic resolution. According to one theoretical estimation, a few thousand images should suffice for calculating the structure of proteins as small as 17 kDa at 3 Å resolution^2^. In practice, however, we are still far away from this theoretical ideal. Thus far, protein complexes that have been successfully reconstructed to high-resolution by single particle analysis (SPA) have molecular weights of ~100 kDa or larger^3^. Here, we report the use of Volta phase plate in determining the structure of human haemoglobin (64 kDa) at 3.2 Å. Our results demonstrate that this method can be applied to complexes that are significantly smaller than those previously studied by conventional defocus-based approaches. Cryo-EM is now close to becoming a fast and cost-effective alternative to crystallography for high-resolution protein structure determination.

---

Given the radiation sensitivity of ice-embedded proteins, the low signal-to-noise ratio (SNR) of cryo-EM images is a limitation for SPA^4^, restricting the size range of proteins that can be studied. In 1995, it was estimated, based solely on physical considerations, that the lower molecular weight limit of single particle cryo-EM would be 38 kDa^5^. These considerations presumed the use of a perfect phase plate. It was suggested that the structure of 100 kDa proteins could be determined at 3 Å resolution from only 10,000 particles. Later, it was proposed that the theoretical limit might be even lower if perfect images could be taken^2, 6^. With the technology at that time and cryo-EM images being far from perfect retaining only 10% of contrast, it seemed that obtaining a 3 Å reconstruction would be reserved for complexes with a molecular weight upwards of 4 MDa^5^. Nowadays, obtaining ~3 Å resolution reconstructions has become almost routine and has been achieved with complexes that are much smaller in size^1^. To date, the smallest protein solved to near-atomic resolution by single particle cryo-EM is the 3.8 Å resolution structure of the 93 kDa isocitrate dehydrogenase^3^. Even so, single particle analysis reconstructions are still strongly biased towards larger symmetric complexes, indicating there is still a long way to go before the full potential of imaging proteins with electrons is reached.

The difficulties in routinely obtaining high-resolution reconstructions of small molecular weight proteins are predominantly owed to poor representation of low spatial frequencies in electron micrographs obtained by conventional transmission electron microscopy (CTEM)^4^. CTEM utilises phase contrast produced by spherical aberration (Cs) and the deliberate defocusing of the microscope’s objective lens. This approach creates oscillations in the contrast transfer function (CTF) of the microscope with some spatial frequencies of the object being transferred poorly, or not at all. One can compensate for this effect by varying the level of defocus from image to image, which is typically in the range of several hundreds to thousands of nanometres. By combining images that have different levels of contrast for given spatial frequencies an accurate representation of an object can be obtained. Nevertheless, the limitations due to reduced SNR resulting from contrast loss remain.

In-focus single particle cryo-EM enabled by the Volta phase plate (VPP) holds the promise of yielding up to a two-fold boost in SNR and therefore enhancing our ability to observe weak phase objects^7^. The SNR of VPP images is high because transfer of contrast of low spatial frequencies is optimal and constant for images taken in focus. Unlike previous phase plate designs, VPP images also retain the high spatial frequencies of the specimen enabling structure determination at near-atomic resolution^8, 9^. However, in-focus imaging with VPP requires very precise focusing^8^ and the typically strong Cs present in cryo-electron microscopes appears to be a limiting factor in attaining resolutions better than 3 Å by in-focus phase plate-imaging^8^.

Enabled by the ability to estimate and correct the phase shift of the VPP in CTFFIND4^10^ and RELION2^11^, we therefore used a hybrid approach combining the strengths of CTEM and VPP^12^ (Fig. 1). This involves applying a defocus of ~500 nm and correcting for the effects of CTF. We opted to apply this strategy to tetrameric Hgb, which mediates oxygen transport in blood and has a molecular weight of 64 kDa and C2 symmetry. We chose Hgb for its iconic status as the first protein structure alongside myoglobin that was solved using X-ray crystallography by Max Perutz in 1960, coincidently by overcoming the phase problem of X-ray crystallography^13^.

**Figure 1.**
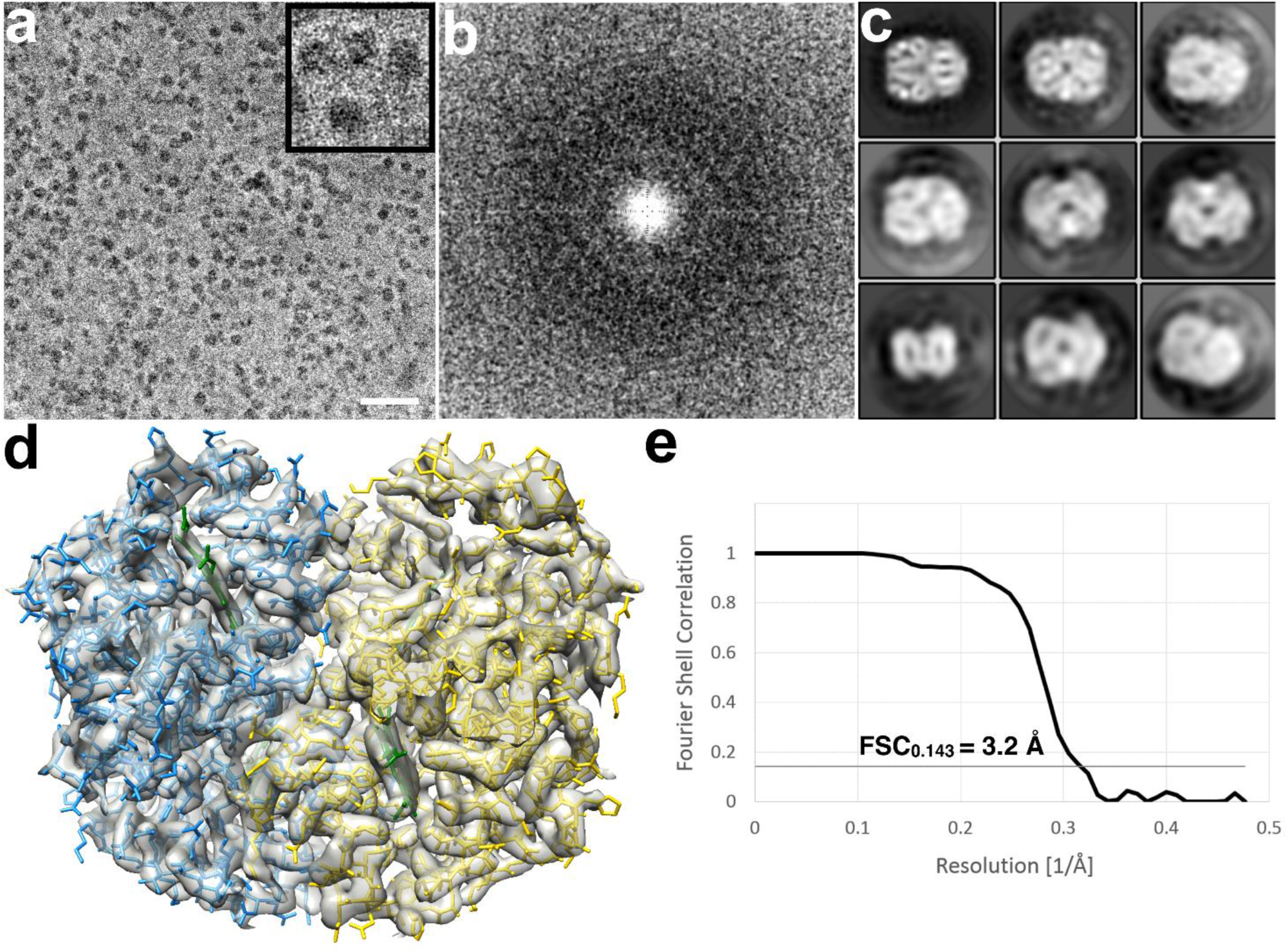
Phase plate-imaging of 64 kDa Hgb. (**a**) Electron micrograph of metHgb recorded at ~500 nm underfocus with the Volta phase plate (VPP) (scale bar = 30nm). (**b**) Power spectrum of the image in (**a**), featuring contrast transfer function (CTF) Thon rings permitting defocus and phase shift estimation. (**c**) 2D class averages of Hgb showing secondary structure elements in projection. (**d**) Reconstructed 3D map and model of Hgb. (**e**) (Gold Standard?) Fourier shell correlation (FSC) plot indicating a resolution of 3.2 Å according to the FSC=0.143 criterion.

Commercially sourced human Hgb is in the non-functional ferric (Fe^3+^) state referred to as metHgb. After vitrification of the metHgb, the sample was subjected to VPP-enabled imaging with multiframe movies taken at low defocus, as described above. The movies were corrected for motion and radiation damage using MotionCor2^14^. Hgb particles were readily discernible in VPP images (Fig. 1a) and could be accurately picked because of their high contrast. 2D classification of automatically picked particles resulted in class averages with recognisable features and striking resemblance to the structure of Hgb (Fig. 1c). Class averages were selected for initial model building in EMAN using the common-line technique and taking advantage of the C2 symmetry^15^. RELION 3D classification and refinement using half-split datasets of particles yielded the final map^16^. The obtained 3D reconstruction had a resolution of 3.2 Å, as determined by the so-called “gold-standard”FSC=0.143 criterion.

At this level of resolution, side-chain densities and prosthetic haem groups are clearly resolved in our reconstruction (Fig. 2a, Fig. S1). We used an MD-based approach for model building and compared our atomic model with three conformers of ferrous (Fe^2+^) Hgb present in a single crystal (PDB 4N7O) adopting the tight (T) and two relaxed states (R1/R2)^17^. Rigid-body fitting was used to dock the α1 subunits yielding a good visual fit with a cross-correlation value of ~69%. Superimposition of docked α1 subunits and corresponding tetramers yields cross-correlation values of 43%, 47% and 62% for T, R1 and R2 states, respectively (Fig. 2b). This observation is in line with the fact that metHgb adopts an R-like state suggesting that conformational states can be determined for small proteins at high-resolution without crystallisation.

**Figure 2.**
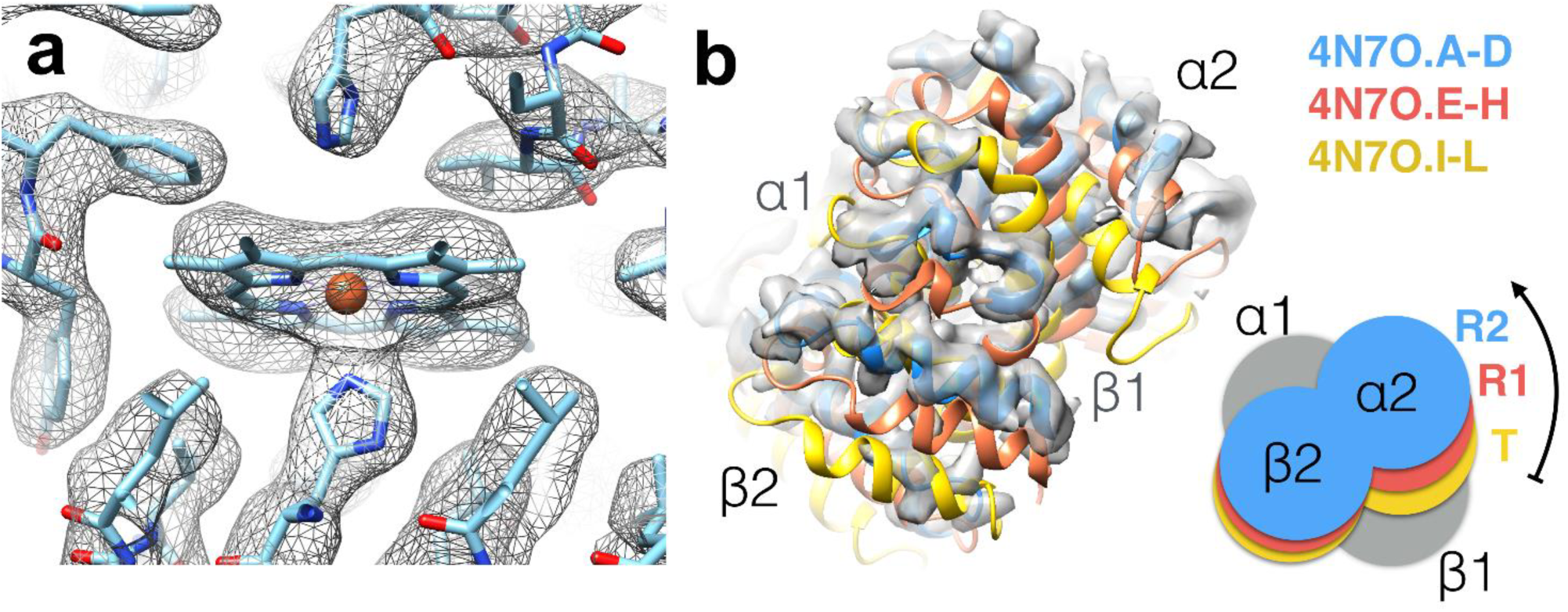
Hgb at 3.2 Å resolution. (**a**) The iron atom of the prosthetic haem group is coordinated by the proximal histidine residue, as evidenced by a strong density connecting them. (**b**) VPP reconstruction fitted with 3 conformers of Hgb present in crystal structure PDB 4N7O. The reconstructed 3D map agrees best with chains A-D of PDB 4N7O representing the R2 state of Hgb.

Our results showcase how cryo-EM can be used to determine which conformational states are present in solution. It has become increasingly clear that simple allosteric models based on discrete states, which are arrested by tight crystal contacts potentially fail to provide a complete structure/function portrait and may be divergent from solution studies^17^. Single particle analysis is inherently better suited than crystallography for visualising the full spectrum of conformational states that proteins adopt^18^. Obtaining high-resolution structures of solution states may indeed be one of the main applications of structure determination by VPP as a technique complementary to X-ray crystallography and nuclear magnetic resonance spectroscopy.

Given the ease of the data acquisition, it can be expected that near-atomic resolution maps will become routine for large parts of the proteome including membrane proteins. In conjunction with improved automation, and next generation direct electron detectors, cryo-EM is likely to become a major player in structure-based small molecule drug discovery for almost any drug target.

## Methods

### Sample preparation

Human Hgb was commercially sourced (Sigma-Aldrich, St. Louis, MO). Frozen-hydrated specimens were prepared on plasma-cleaned Quantifoil R1.2/1.3 holey carbon EM grids (Quantifoil, Großlöbichau,Germany) using a Vitrobot Mark III (FEI, Hillsboro, OR) 5 s blotting time, 85% humidity and −5 mm blotting offset.

### Data acquisition

Automated data collection was performed on a Titan Krios electron microscope (FEI, Hillsboro, OR) operated at 300 kV and equipped with a K2 Summit direct detector, a Quantum energy filter (Gatan, Pleasanton, CA) and an FEI Volta phase plate (FEI, Hillsboro, OR) using SerialEM software. Movies comprising 40 frames and a total dose of 40 e^−^per Å^2^ were recorded on a K2 Summit direct detection camera (Gatan) at a calibrated magnification of 95,200 corresponding to a magnified pixel size of 0.525 Å.

### Data processing

The recorded movies were subjected to motion correction with MotionCor2^14^. Particles were picked with Gautomatch (developed by Zhang K, MRC Laboratory of Molecular Biology, Cambridge, UK, http://www.mrc-lmb.cam.ac.uk/kzhang/Gautomatch/). Subsequently, the 283,600 picked particles were extracted in Relion2 using a box size of 100 pixels^11^. After performing 2D classification in Relion2, the best-looking 2D class averages were selected to build an initial model in EMAN using the common-lines approach^15^. After two rounds of 3D classification 164,300 particles from the best looking class were subjected to 3D auto-refinement in Relion2. The final map was sharpened using a negative B-factor of 200. Local resolution was calculated with *blocres* from the Bsoft package^19^. Flexible fitting of the Hgb crystal structure was performed using the NAMD routine in MDFF^20^ followed by real-space refinement in PHENIX^21^. The data collection, refinement parameters and model statistics are summarised in Table 1.

**Table 1.**
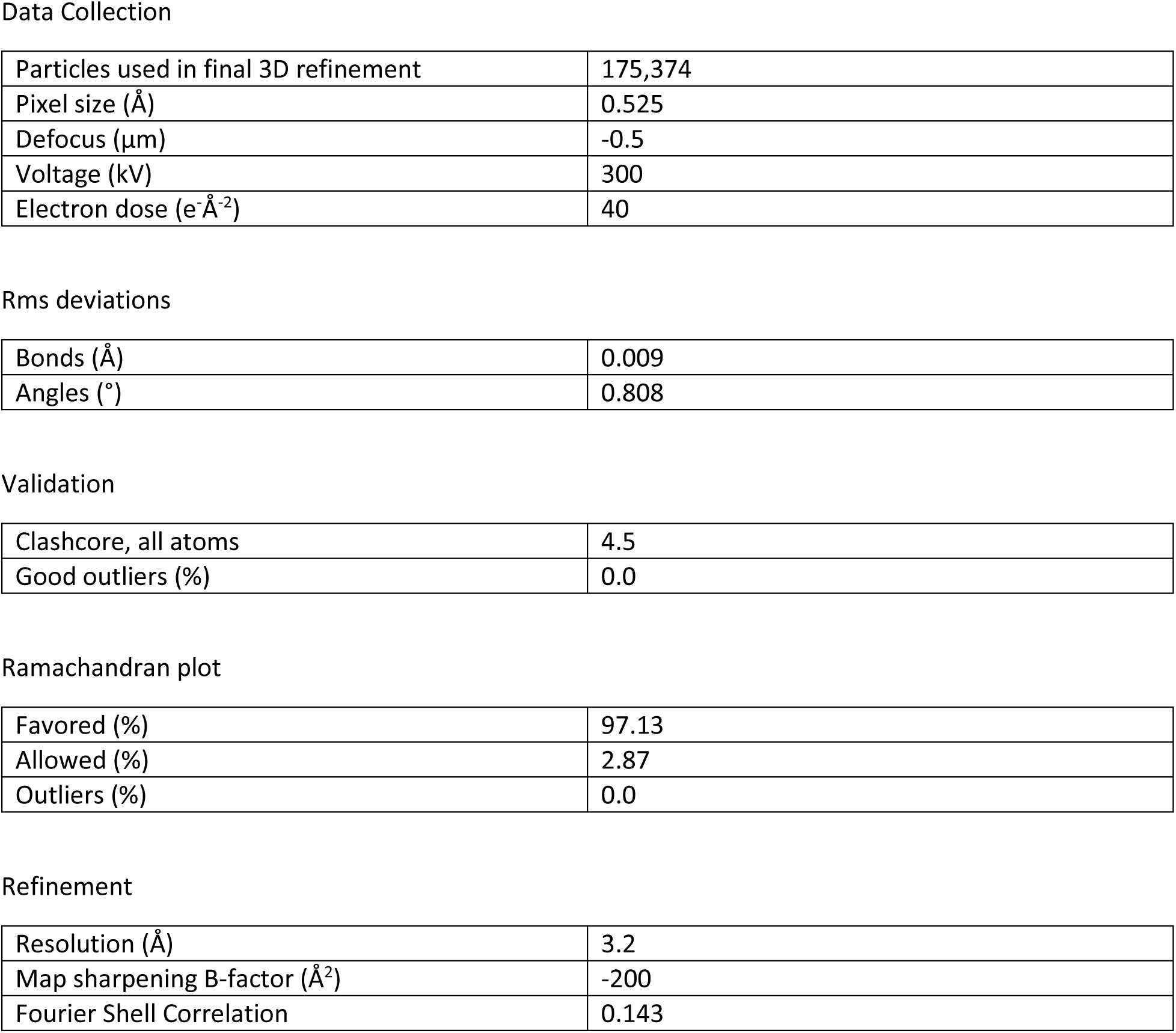

Data Collection
|  |  |
| --- | --- |
| Particles used in final 3D refinement | 175,374 |
| Pixel size (Å) | 0.525 |
| Defocus (μm) | -0.5 |
| Voltage (kV) | 300 |
| Electron dose (e<sup>-</sup> Å<sup>-2</sup>) | 40 |
| Rms deviations |  |
| Bonds (Å) | 0.009 |
| Angles (°) | 0.808 |
| Validation |  |
| Clashcore, all atoms | 4.5 |
| Good outliers (%) | 0.0 |
| Ramachandran plot |  |
| Favored (%) | 97.13 |
| Allowed (%) | 2.87 |
| Outliers (%) | 0.0 |
| Refinement |  |
| Resolution (Å) | 3.2 |
| Map sharpening B-factor (Å<sup>2</sup>) | -200 |
| Fourier Shell Correlation | 0.143 |

## Accession codes

The cryo-EM map and atomic coordinates of Hgb were deposited to the Electron Microscopy Data Bank (EMDB) and Protein Data Bank (PDB) under accession codes EMD-3488 and PDB-5ME2, respectively. Raw data was made available at the Electron Microscopy Pilot Image Archive (EMPIAR) with accession code EMPIAR-10105.

## Acknowledgments

We thank Prof. Jürgen Plitzko for his technical support. We also acknowledge Dr. Alexis Rohou and Sjors Scheres for support with CTFFIND4 and Relion2, respectively. We thank Dr. Matthew Belousoff for model building of the Hgb atomic model. We also thank Dr. Mike Strauss and Dr. Shelley Robison for critical reading of the manuscript. This work was supported by the Multi-modal Australian ScienceS Imaging and Visualisation Environment (www.massive.org.au).

## Author contributions

MK, MR and RD were responsible for the conception, design, data analysis and interpretation of experiments. MK performed sample preparation and imaging. All authors wrote the manuscript.

---

**Figure S1.**
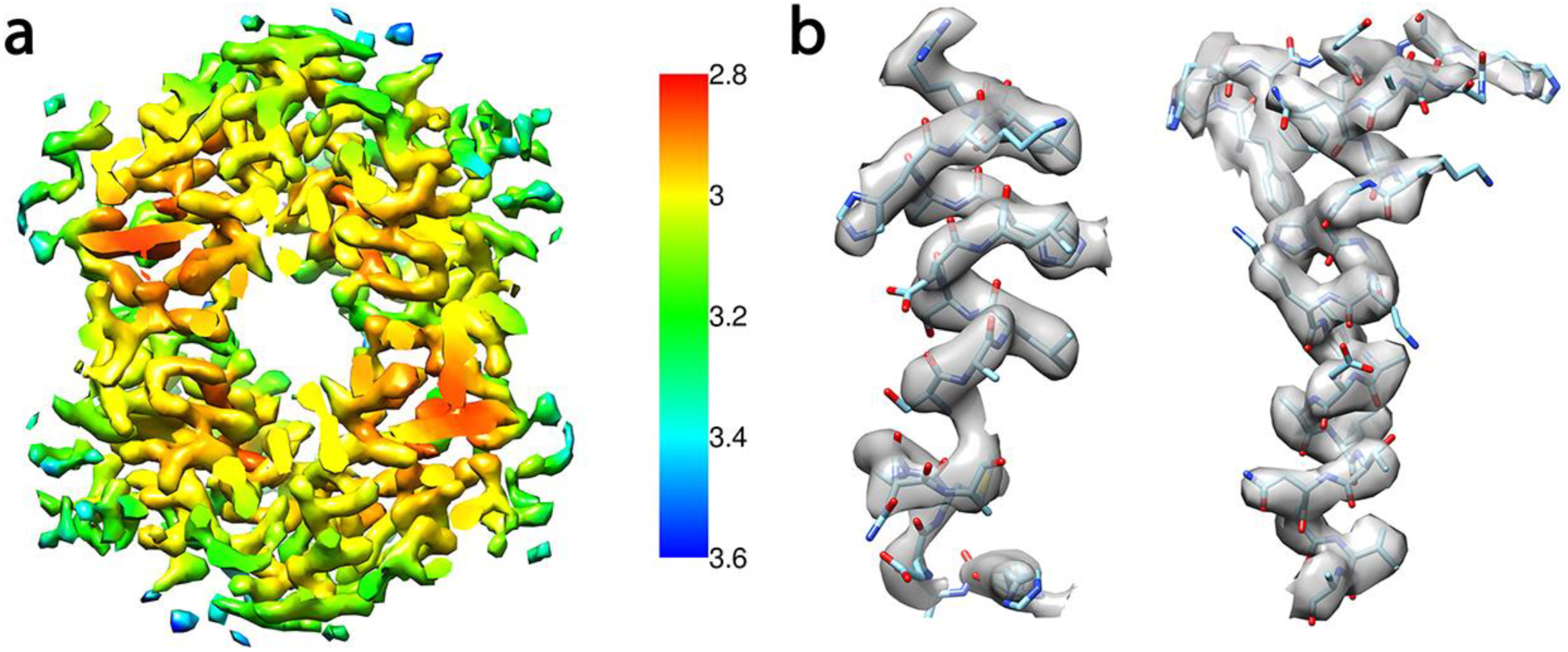
(**a**) Local resolution estimation. (**b**) Magnified details of the 3D map with refined model.

